# Graded Visual Consciousness During the Attentional Blink

**DOI:** 10.1101/2021.01.15.426792

**Authors:** Anna Eiserbeck, Alexander Enge, Milena Rabovsky, Rasha Abdel Rahman

## Abstract

One of the ongoing debates about visual consciousness is whether it can be considered as an all-or-none or a graded phenomenon. This may depend on the experimental paradigm and the task used to investigate this question. The present event-related potential study (*N* = 32) focuses on the attentional blink paradigm for which so far only little and mixed evidence is available. Detection of T2 face targets during the attentional blink was assessed via an objective accuracy measure (reporting the faces’ gender), subjective visibility on a perceptual awareness scale (PAS) as well as event-related potentials time-locked to T2 onset (components P1, N1, N2, and P3). The behavioral results indicate a graded rather than an all-or-none pattern of visual awareness. Corresponding graded differences in the N1, N2, and P3 components were observed for the comparison of visibility levels. These findings suggest that conscious perception during the attentional blink can occur in a graded fashion.

## Introduction

A popular analogy in regard to consciousness is that of light. The metaphor of an on/off switch has been used, for example, in relation to anatomical structures considered to be important for the presence of consciousness (e.g., Blumenfeld, 2014). But does the implied idea of a dichotomous concept of conscious perception, where there is either light or darkness, really correspond to the underlying structure of the phenomenon? Or would it perhaps be more accurate to compare it to, say, a light dimmer that allows for continuously varying degrees of brightness? In a simplified manner, this analogy represents a question that has increasingly been debated in consciousness research—and in particular the field of visual consciousness—in recent years: What is the nature of conscious perception? Can it be understood as an all-or-nothing phenomenon where something is either fully perceived or not perceived at all, or do gradations in visual consciousness exist?

After this debate has received a surge of attention through articles by Sergent and Dehaene (2004) as well as Ramsoy and Overgaard (2004), there now exists considerable evidence in both directions. Since this has already been descrid in detail elsewhere (e.g. Fazekas & Overgaard, 2018; Förster et al., 2020; Jonkisz et al., 2017; Kiefer & Kammer, 2017; Windey et al., 2014), we will refrain from an extensive review of the evidence in general. As suggested previously (Pretorius, 2014), the question may not be whether graded states of consciousness exist at all, but under which circumstances they can be observed. Among other factors, this may depend on the degradation technique used to investigate visual consciousness (Experiment 1 in Pretorius, 2014). This article focuses on the attentional blink paradigm—an often-used method in the study of consciousness—for which, as yet, scarce evidence regarding this matter is available.

The attentional blink describes the phenomenon that in a rapid serial visual presentation (RSVP) task, the detection of the second (T2) of two target stimuli presented in a stream of distractor stimuli is often impaired if it follows the first target stimulus (T1) within a period of 200 to 500 ms as compared to a longer (or shorter) interval between both targets (Raymond et al., 1992). As yet, the mechanisms behind this phenomenon are not fully resolved but in general terms explained by a disruption of attentional processes and/or of encoding in working memory due to the ongoing processing of T1 (for a recent review, see Zivony & Lamy, 2020). Often in investigations of the attentional blink only the contrast between trials in which T2 was “seen” versus “not seen” (or “detected” versus “not detected”) was considered, which implicitly contains the assumption of an all-or-none pattern of conscious perception. Even the term *attentional blink* itself seems to imply a dichotomous separation of either seeing, or not seeing when blinking. However, some studies have also specifically addressed the question of the form of consciousness during the attentional blink and in doing so have come to differing conclusions.

### Behavioral Evidence in Favor of and Against Dichotomous Consciousness During Attentional Blink

In a seminal study investigating the form of consciousness during the attentional blink, Sergent and Dehaene (2004) assessed the detection of T2 number words using a continuous visibility scale with 21 contiguous positions. The response distribution obtained during the attentional blink was focused on the ends of the scale and represented a mixture of the distribution of “seen” states (as obtained when T2 was presented in a single task condition) and “unseen” states (as obtained when T2 was absent in a single task condition), thus representing an all-on-none pattern of conscious perception. Using the same study design, this finding has also been replicated and extended by the investigation of underlying neural correlates (Sergent et al., 2005), as outlined below. Further evidence for dichotomous conscious perception during the attentional blink comes from a study by Asplund et al. (2014) comparing the precision of T2 identification between lags. The experiments are based on the assumption that graded awareness should be reflected in an increasingly precise perception of T2 with longer T1–T2 lags. In two experiments—the first one investigating T2 color identification, the second one investigating T2 face identification—participants selected T2 stimulus identity on a continuous response wheel. Differences between lags were found in the *report probability* of T2, i.e., whether the response is in proximity to the correct color value / face identity or not, but not in the *precision* with which the color / face stimulus was identified, i.e., how close exactly the given response is to the correct one. The authors concluded that in the attentional blink, conscious perception occurs in an all-or-none manner without differences in precision.

Using different experimental designs and measures (a subjective visibility rating versus an objective accuracy measure) Sergent and Dehaene (2004), and Asplund et al. (2014), thus, reached the same conclusion in favor of a dichotomous pattern of conscious perception during the attentional blink. However, in both cases the observed all-or-none pattern may be the result of specific experimental parameters. Regarding the study by Sergent and Dehaene (2004), a “two-stage” process underlying the evaluation of T2 word visibility may have biased the responses towards a dichotomous pattern: In order to be able to judge the word’s visibility, it is essential to first identify the array of characters as a word (in contrast to the random arrangement of letters used as distractors). If there were cases of partial or degraded perception of the stimulus, participants might not have been able to do so and in these cases reported “not seen” even though they had a partial/vague visual impression of the stimulus (Elliott et al., 2016; also see Stein & Peelen, 2020, for evidence and discussion on the importance of distinguishing between stimulus detection and discrimination). The opposite direction is also conceivable: Seeing only parts of the word may have been sufficient for being able to identify the word and thus report a high visibility (Nieuwenhuis & de Kleijn, 2011). Furthermore, with a fine-grained scale such as the 21-point scale employed by Sergent and Dehaene (2004) it may be difficult to differentiate between the many intermediate levels, which may result in a stronger focus on the ends of the scale, especially since only the endpoints were labelled (see Overgaard et al., 2006). Further evidence (Pretorius et al., 2016) showed that, even though the 21-point scale is generally suitable to capture graded states of consciousness, in doing so it may be less sensitive compared to scales with less response options (a four-, seven-, or three-point scale), as indicated by a lower proportion of responses in an intermediate range as compared to the other scales when a direct comparison between scales was enabled through transformation. Regarding an objective accuracy measure of T2 detection as used in Asplund et al. (2014), it needs to be noted that, even if the precision with which a stimulus is identified exhibits an all-or-none pattern, this does not preclude the possibility that graded differences with respect to other dimensions—such as the perceived intensity and/or temporal stability of the percept—may be found (see Fazekas & Overgaard, 2018). In line with this idea, evidence indicates that T2 accuracy and visibility (i.e., subjective experience of T2) are distinguishable (Pincham et al., 2016). Therefore, the precision with which a certain task regarding the stimulus can be solved does not necessarily equal the overall perceived visibility or quality.

Evidence for an influence of the experimental design on the obtained pattern of results as well as initial evidence for graded consciousness during the attention blink was presented by Nieuwenhuis and de Kleijn (2011) who varied the type of target stimulus and the measure with which conscious perception was assessed. In a first experiment with a design similar to that of Sergent and Dehaene (2004), in which words were used as T2 stimuli in combination with a 7-point visibility scale (instead of a 21-point version as in Sergent & Dehaene, 2004), a dichotomous pattern (more precisely described as “discontinuous transition”) was replicated. However, when individual number characters had to be identified with the same rating scale, participants used the scale in a continuous fashion, indicating graded changes in conscious perception.^1^ In a study using the four-point perceptual awareness scale (PAS; Ramsøy & Overgaard, 2004), Pretorius (2014) investigated visual consciousness for words versus shapes in an attentional blink task and a visual masking condition. Graded states of visual consciousness were observed under all conditions. Yet, noticeably, the particular combination of the attentional blink paradigm and word stimuli (as opposed to shape stimuli and/or the masking technique) yielded the lowest amount of graded awareness reports (i.e., intermediate ratings on the scale). These studies corroborate the idea that the dichotomous pattern observed in Sergent and Dehaene (2004) may be due to specific experimental parameters and that a graded pattern of visibility reports can be obtained with variations of the stimulus material and visibility scale.

### Evidence From Event-Related Potential Studies

In the present study we employ event-related brain potentials (ERPs) to investigate how the perceived visibility of T2 stimuli during the attentional blink is reflected in components associated with perception and consciousness (for reviews, see Förster et al., 2020; Koivisto & Revonsuo, 2010). Although the question of the neural correlates of consciousness has not yet been resolved, in ERP research the focus has narrowed down especially on two components: The visual awareness negativity (VAN) represents a relative negativity for seen as compared to unseen stimuli at posterior sites spanning the time range of the N1 and N2 components, with a peak latency around 200 to 250 ms after stimulus onset. The late positivity (LP) is a relative positivity in the time range of the P3b, peaking approximately 400 ms after stimulus onset. Although differences in the LP/P3b time range have frequently been observed when comparing detected and undetected stimuli across various paradigms, increasing evidence indicates that this component may not be a marker of conscious perception itself, but rather of the *subjective report* of stimulus detection, thus reflecting post-perceptual processing (for a review, see Koch et al., 2016). For example, it has been found that consciously perceived but task-irrelevant/non-reported stimuli do not elicit a P3b (Cohen et al., 2020; Pitts et al., 2014). Instead, reviews across different paradigms consider the VAN as the earliest and most consistent correlate of conscious perception (in terms of phenomenal consciousness, i.e., the subjective experience of seeing; Förster et al., 2020; Koivisto & Revonsuo, 2010).

As yet, the only attentional blink study which investigated ERP differences between different levels of visibility is the one by Sergent et al. (2005).^2^ Using the same experimental design as Sergent and Dehaene (2004), the dichotomous distribution of visibility ratings was replicated, and this was accompanied by a dichotomous activation pattern of the P3b (as well as the N3 und P3a). Interestingly, a gradual pattern was found during the N2 time range as the earliest component for which differences between visibility levels were observed. However, based on the bimodal distribution of visibility ratings as well as theoretical considerations (the global neuronal workspace model; Dehaene et al., 2003), the authors concluded that access to consciousness during the attentional blink occurs in an all-or-none fashion and may depend on the optional triggering of later components displaying a bimodal pattern of activity. The graded activity observed in the N2 was taken to reflect a pre-conscious processing stage, i.e., in this view, pre-conscious processing may be graded, but the access to consciousness occurs in an all-or-none fashion. This interpretation is challenged by new evidence from consciousness studies on the whole as given above suggesting that activity during the N2/VAN time range may be a better indicator of conscious perception than the P3b. In any case, the observation that the dichotomous rating pattern obtained with the particular study design does not necessarily generalize to all attentional blink studies raises the need to further investigate this matter using different study designs.

### Current Study

In summary, there exists behavioral evidence for both all-or-none and graded visual consciousness during the attentional blink. ERP evidence regarding this matter comes from only one study (Sergent et al., 2005) and may not generalize to attentional blink studies with different designs. This indicates the necessity to investigate this matter again under different experimental conditions, which is the aim of this study.

The current study investigates the form of conscious perception of T2 face stimuli in the attentional blink with behavioral measures and ERPs obtained from the EEG. Behavioral measures consisted of an objective accuracy measure (reporting the faces’ gender) as well as subjective visibility judgments. A variant of the PAS scale was used which has been developed to be sensitive for graded differences in conscious visual perception (Ramsøy & Overgaard, 2004). If conscious perception during the attentional blink occurs in a graded manner, this should be reflected in the presence of intermediate responses on the visibility scale (rather than ratings being clustered at end points). Linking subjective ratings with accuracy reports as well as reaction times allowed to examine systematic differences between levels of perceived visibility. Thereby, higher visibility ratings should go along with a higher accuracy and faster reaction times in the objective task (Andersen et al., 2016). In the ERPs, we compared activity for the four visibility levels in components on which consciousness research has mainly focused (Koivisto & Revonsuo, 2010; Sergent et al., 2005): P1; N1, and N2 (as part of the VAN); and P3b (/LP).

## Methods

### Participants

A re-analysis of datasets from an attentional blink study investigating influences on the visual consciousness of faces was conducted (Eiserbeck et al., in prep.): The sample comprised 32 native German speakers (21 female) with a mean age of 26.1 (*SD* = 6.65) years and with normal or corrected-to-normal vision. Participants had provided written informed consent prior to participation. The study was conducted according to the principles expressed in the Declaration of Helsinki and was approved by the local Ethics Committee. Participants received either course credit or monetary compensation of C8 per hour.

### Materials

Three types of images were used in the RSVP stream:

- T1 target stimuli consisted of 36 images displaying either the face of a dog or a similarly looking blueberry muffin, all converted to greyscale and cropped to the same oval shape.
- T2 target stimuli consisted of 24 portraits of faces (12 female) with Caucasian appearance, displaying neutral emotional expressions, taken from the Chicago Face Database (CFD; Ma et al., 2015). The pictures were converted to greyscale and cropped so that no hair and ears were visible. Histograms (i.e., the distributions of brightness values) of the images were equated using the SHINE toolbox (Willenbockel et al., 2010) in MATLAB (Version R2016a).
- As distractor images 12 abstract stimuli—visually similar to T2 targets—were used (see Müsch et al., 2012, for the importance of target-distractor-similarity). For this purpose, 12 additional faces from the CFD were processed in the same way as described above and their facial features were rotated and displaced in different positions.

Stimuli were presented on a 19-inch LCD monitor with a 75-Hz refresh rate. They were displayed on a grey background with a size subtending 5.8° vertical visual angle and 4.3° horizontal visual angle (viewing distance: 70 cm).

### Procedure

For each trial of the attentional blink task, first a fixation cross was presented for 500 ms. Then, 13 pictures were shown in rapid succession, with a presentation time of 107 ms each and without a time interval between pictures (for a graphical illustration of the trial structure, see Figure 1A). Regular trials contained 11 distractor images, which were presented in randomized order, and two targets: a dog or muffin (T1) and a face (T2). T2 (if present) was always presented as the 10th stimulus whereas T1 position varied: It was either presented as the third stimulus (entailing a lag of seven items between T1 and T2; long lag) or as the seventh stimulus (entailing a lag of three items; short lag). The task comprised 696 trials in total. As T1, in 50 % of cases a dog was shown and in 50 % a muffin. Within each lag, T2 was absent in 60 trials (17 %) and instead another distractor was presented. Trial types (short or long lag, T2 present or absent) were presented in randomized order.

**Fig. 1.**
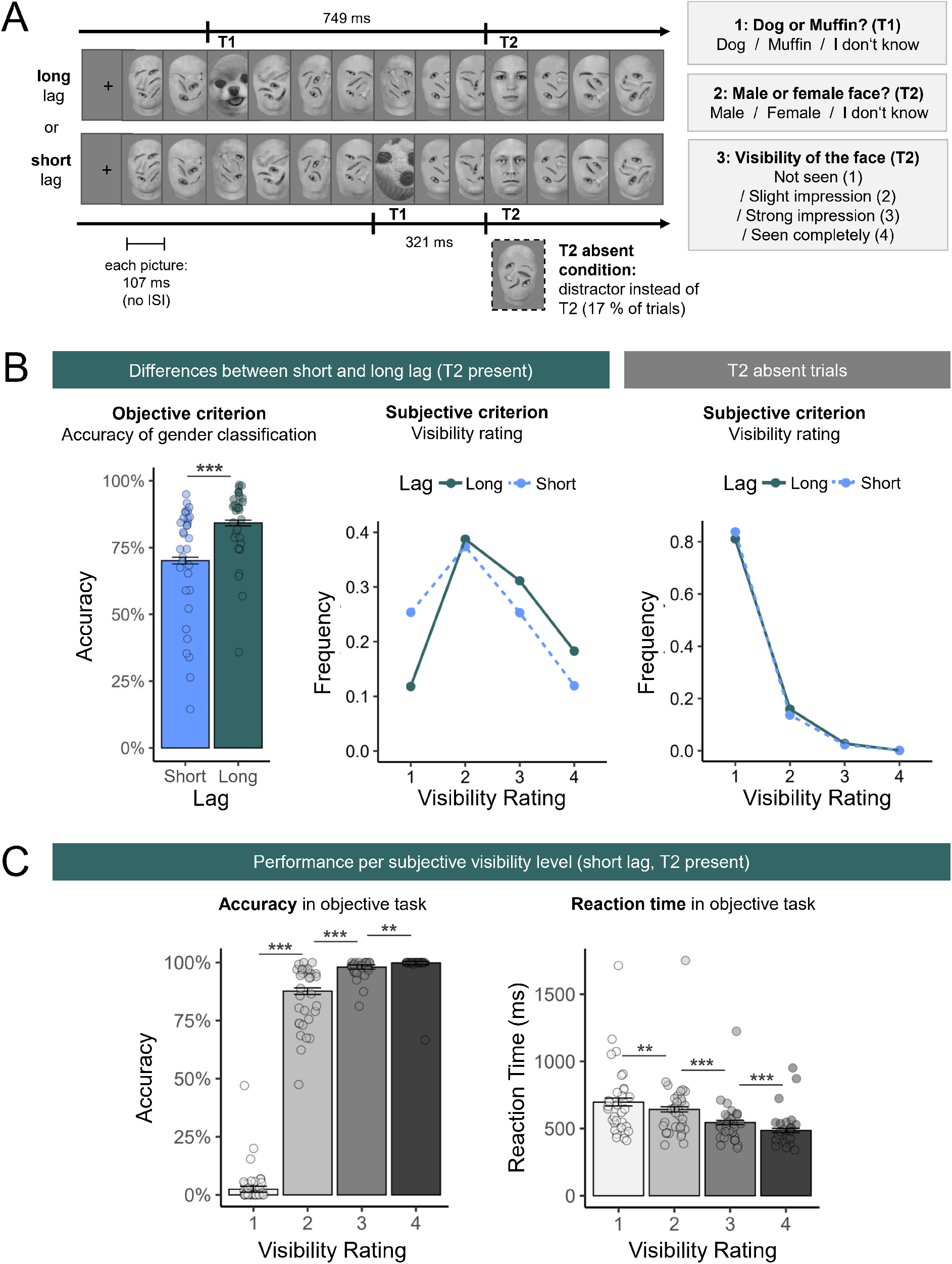
Design of the attentional blink task and behavioral results. **(A)** Structure of long and short lag trials. After each trial, participants answered three questions regarding T1 identity, T2 gender, and T2 visibility. **(B)** Results for objective criterion (gender classification task) and subjective criterion (visibility ratings) per lag in T2 present and T2 absent trials. Error bars depict 95% confidence intervals; circles in the bar plot represent means for individual participants. For illustration of corresponding by-item means, see Figure A2. **(C)** Accuracy and reaction time in objective task as a function of visibility rating. Please note that the very low mean accuracy rate for *not seen* is due to the fact that participants were explicitly instructed not to guess and instead to choose the response option “I don’t know” if they were unsure about the face’s gender. Error bars depict 95% confidence intervals; circles in bar plots represent means of individual participants. For illustration of corresponding by-item means, see Figure A2. Significance codes: *** *p* < .001, ** *p* < .01. Visibility levels: 1 = *not seen*, 2 = *slight impression*, 3 = *strong impression*, 4 = *seen completely*.

Participants were instructed to look for the dog or muffin and the face. They were informed that both targets are equally important, but that not every sequence contains a face. After each trial, participants indicated via response keys (a) whether they saw the image of a dog or a muffin as T1 (options: *dog / muffin / I don’t know*), (b) whether they saw a male or a female face as T2 (options: *male / female / I don’t know*), and (c) how clear their subjective impression of T2 was on a four-point perception awareness scale (PAS; Ramsøy & Overgaard, 2004; options: *not seen / slight impression / strong impression / seen completely*). There were no time constraints for answering.

The attentional blink task described here was part of an experiment which comprised additional parts: pre-ratings of facial trustworthiness and facial expression, a learning phase in which emotional knowledge about the T2 faces was obtained before the attentional blink task, and post-ratings of facial trustworthiness and facial expression. For the purpose of the present study, only the procedure of the attentional blink task is described, since the other tasks are not directly relevant for the research question at hand.

### EEG Recording and Processing

The EEG was recorded with Ag/AgCl electrodes at 62 scalp sites according to the extended 10–20 system at a sampling rate of 500 Hz and with all electrodes referenced to the left mastoid. An external electrode below the left eye was used to measure electrooculograms. During recording, low- and high-cut-off filters (0.016 Hz and 1000 Hz) were applied and all electrode impedances were being kept below 10 kΩ. After the experiment, a calibration procedure was used to obtain prototypical eye movements for later artifact correction. Processing and analyses of the data were based on the EEG-processing pipeline by Frömer, Maier, and Abdel Rahman (2018). An offline pre-processing was conducted using MATLAB (Version R2016a) and the EEGLAB toolbox (Version 13.5.4b; Delorme & Makeig, 2004). The continuous EEG was rereferenced to a common average reference and eye movement artifacts were removed using a spatio-temporal dipole modeling procedure with the BESA software (Ille et al., 2002). The corrected data were low-pass filtered with an upper passband edge at 40 Hz. Subsequently, they were segmented into epochs of −200 to 1,000 ms relative to T2 onset, and baseline-corrected using the 200 ms pre-stimulus interval. Segments containing artifacts (absolute amplitudes over ±150 µV or amplitudes changing by more than 50 µV between samples) were excluded from further analysis.

## Data Analyses and Results

All analyses were conducted in R (Version 3.5.1; R Core Team, 2014) using the lme4 package (Version 1.1-23; Bates, et al., 2015) and the lmerTest package (Version 3.0-1; Kuznetsova et al., 2017) to calculate p-values via Satterthwaite approximation in case of linear mixed models (while in case of generalized linear mixed models, p-values were based on the Wald z-test implemented in lme4). In all analyses we aimed to include the maximal random effects structure which still enables model convergence and does not lead to over-fitting. If necessary, random effects were excluded based on least explained variance (using the rePCA function; see Bates et al., 2018).

### Behavioral Data

#### Attentional Blink Effect

Analyses investigating the effect of *lag* (short / long) on objective performance and subjective visibility were conducted to verify a functioning attentional blink paradigm with the typical pattern of attenuated detection in short as compared to long lag trials. To this end, two separate models were utilized. In a generalized linear mixed model, the dependent variable *accuracy* (with the binary outcome of 1 or 0, relating to “correct gender categorization” and “missed/incorrect gender categorization”) was predicted by the single fixed effect *lag* (short / long; effect-coded as −0.5, 0.5), taking into account random intercepts for participants and items, as well as by-participant and by-item random slopes for *lag*. The analysis yielded a main effect of *lag* (*b* = 0.98, *z* = 5.99, *p* < .001), as reflected in a higher mean accuracy in long (84%) as compared to short lag trials (70%; see Figure 1B for graphical display and Table A1 for model output).

A second linear mixed model was used to investigate the effect of *lag* (short / long; effect-coded as −0.5, 0.5), on visibility rating (implemented as numeric values from 1 to 4) in T2-present trials, taking into account random intercepts for participants and items, as well as by-participant random slopes for *lag*.^3^ The analyses yielded a main effect of *lag* (*b* = 0.32, *t*(31.00) = 4.67, *p* < .001), as reflected in a higher mean visibility rating in long (2.56 points) as compared to short lag trials (2.24 points; see Table A1 for model output).

#### Distribution of Visibility Ratings

The distribution of visibility ratings for both short and long lag conditions can be examined in Figure 1B (for both T2 present as well as T2 absent trials); the corresponding percentage rates are displayed in Table 1. As apparent, the T2 present short lag (as well as the long lag) condition is characterized by a high ratio of intermediate scale responses while the distribution in T2 absent trials is focused on the low end of the scale.

**Table 1.** Distribution of visibility ratings (relative frequency) per condition

| Visibility rating | T2 present |  | T2 absent |  |
| --- | --- | --- | --- | --- |
|  | Short lag | Long lag | Short lag | Long lag |
| Not seen (1) | 25.62 % | 11.99 % | 83.81 % | 81.05 % |
| Slight impression (2) | 37.86 % | 39.51 % | 13.73 % | 15.94 % |
| Strong Impression (3) | 24.89 % | 30.70 % | 2.29 % | 2.83 % |
| Seen completely (4) | 11.63 % | 17.80 % | 0.17 % | 0.18 % |

#### Accuracy per Visibility Level

Additional generalized linear mixed model analyses were conducted to investigate the connection of the objective and the subjective criterion in short lag trials. To this end, *accuracy* was predicted by the fixed effect *visibility* (coded as a factor with four levels), taking into account random intercepts of participants and items. Sliding difference contrast coding was applied, which enables testing adjacent factor levels against each other. The analyses revealed significant differences in accuracy for the comparison of all adjacent factor levels (with higher accuracy for the higher visibility rating; all *p*s ≤ .001); for statistical output see Table A2. A graphical display of accuracy rates per visibility rating level can be found in Figure 1C; corresponding means and 95% confidence intervals (CIs) are displayed in Table 2.

**Table 2.**
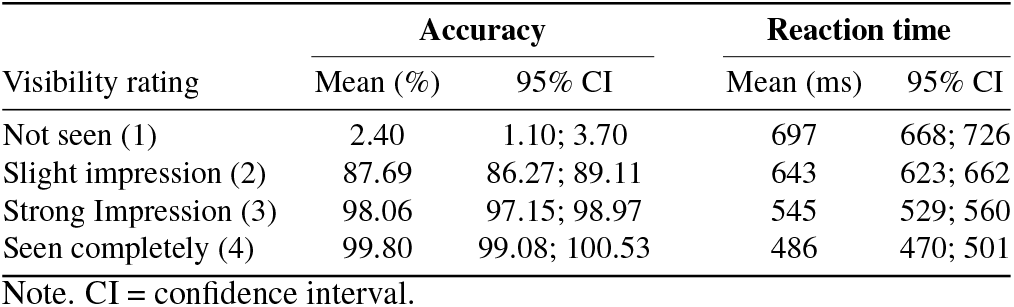
Accuracy and reaction times in objective task per visibility level in T2 present short lag trials

| Visibility rating | Accuracy |  | Reaction time |  |
| --- | --- | --- | --- | --- |
|  | Mean (%) | 95% CI | Mean (ms) | 95% CI |
| Not seen (1) | 2.40 | 1.10; 3.70 | 697 | 668; 726 |
| Slight impression (2) | 87.69 | 86.27; 89.11 | 643 | 623; 662 |
| Strong Impression (3) | 98.06 | 97.15; 98.97 | 545 | 529; 560 |
| Seen completely (4) | 99.80 | 99.08; 100.53 | 486 | 470; 501 |

Note. CI = confidence interval.

#### Reaction Time per Visibility Level

Further linear mixed model analyses were conducted to investigate the connection of visibility rating and reaction times for gender classification in short lag trials. To this end, reaction time was predicted by the fixed effect *visibility*, taking into account random intercepts of participants and items. As in the previous model, sliding difference contrast coding was applied. To better meet the assumption of normality for linear mixed model analyses, reaction time data was trimmed, excluding trials with reactions times faster than 200ms and slower than 2.5 standard deviations above the mean of individual participants, and log-transformation was applied. Data trimming led to an exclusion of 1010 of 8540 short lag T2-present trials (11.83%; mean excluded trials per participant = 32, *SD* = 39). The analyses revealed significant differences in reaction times for the comparison of all adjacent factor levels (with reaction being faster the higher the rated visibility; all *p*s *≤*.006); for statistical output see Table A2. A graphical display of (trimmed, untransformed) reaction times per visibility rating level can be found in Figure 1C; corresponding means and 95% CIs are displayed in Table 2.

#### ERPs

Time windows and electrode sites taken into account for each component were based on previous research, as well as visual inspection of the activity for T2 present versus T2 absent trials to determine the exact time range for analyses (i.e., the factor visibility itself was thereby not taken into account). Analyses comprised the following time ranges and electrodes: P1 from 90–130 ms at electrodes Oz, O1, O2, POz, PO3, PO4, PO7, PO8 (see Maier & Abdel Rahman, 2018), N1/N170 from 150–220 ms at electrodes TP9, TP10, P7, P8, PO9, PO10, O1, O2 (see Hinojosa et al., 2015; Itier & Taylor, 2004), N2 from 220–300 ms at electrodes TP9, TP10, P7, P8, PO9, PO10, O1, O2 (see Del Cul et al., 2007; Sergent et al., 2005), and P3b from 400–800 ms at electrodes Pz, CPz, CP1, CP2, Cz, CP3, CP4 (see Cohen et al., 2020; Sergent et al., 2005).

Single-trial ERPs were averaged across the respective time window and electrodes. They were analyzed with linear mixes models (LMMs) including random intercepts for subjects and items, with mean amplitude of the respective component serving as dependent variable. In order to account for activity caused by the RSVP stream (rather than the T2 stimulus), for each component, the mean amplitude value across participants during T2 absent short lag trials was subtracted as a constant from the dependent variable. This made it possible to obtain more plausibly interpretable estimates for the intercept, but otherwise does not change the results in any way. The models included the single fixed factor *visibility* (4 levels: not seen / slight impression / strong impression / seen completely). Sliding difference contrast coding was applied in order to test adjacent factor levels against each other. ERP analyses focused on short lag trials, which represent the condition of reduced attention relevant to the hypotheses.

Table 3 contains estimates (regression coefficients *b*) of the fixed effects, confidence intervals, *t*-values and *p*-values for the analysis of short lag trials. Graphical illustration and to-pographic displays can be found in Figure 2.

**Table 3.**
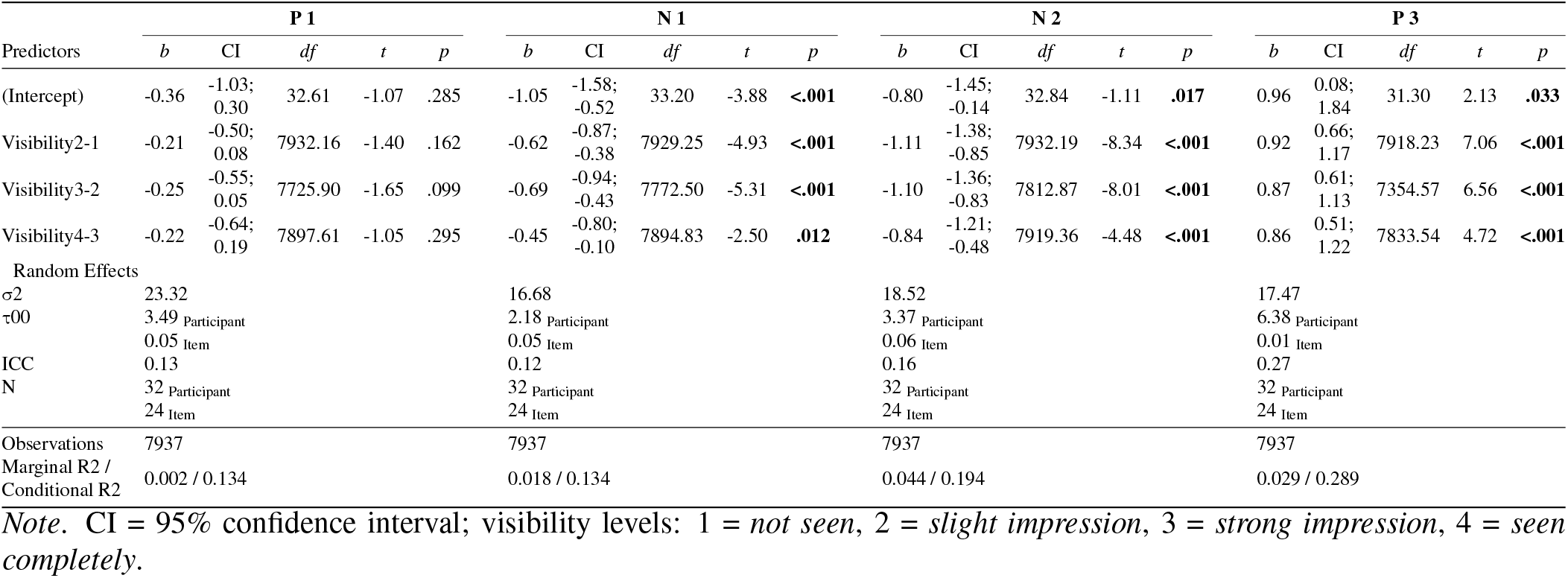
LMM statistics for ERP analyses of T2 present short lag trials.

**Fig. 2.**
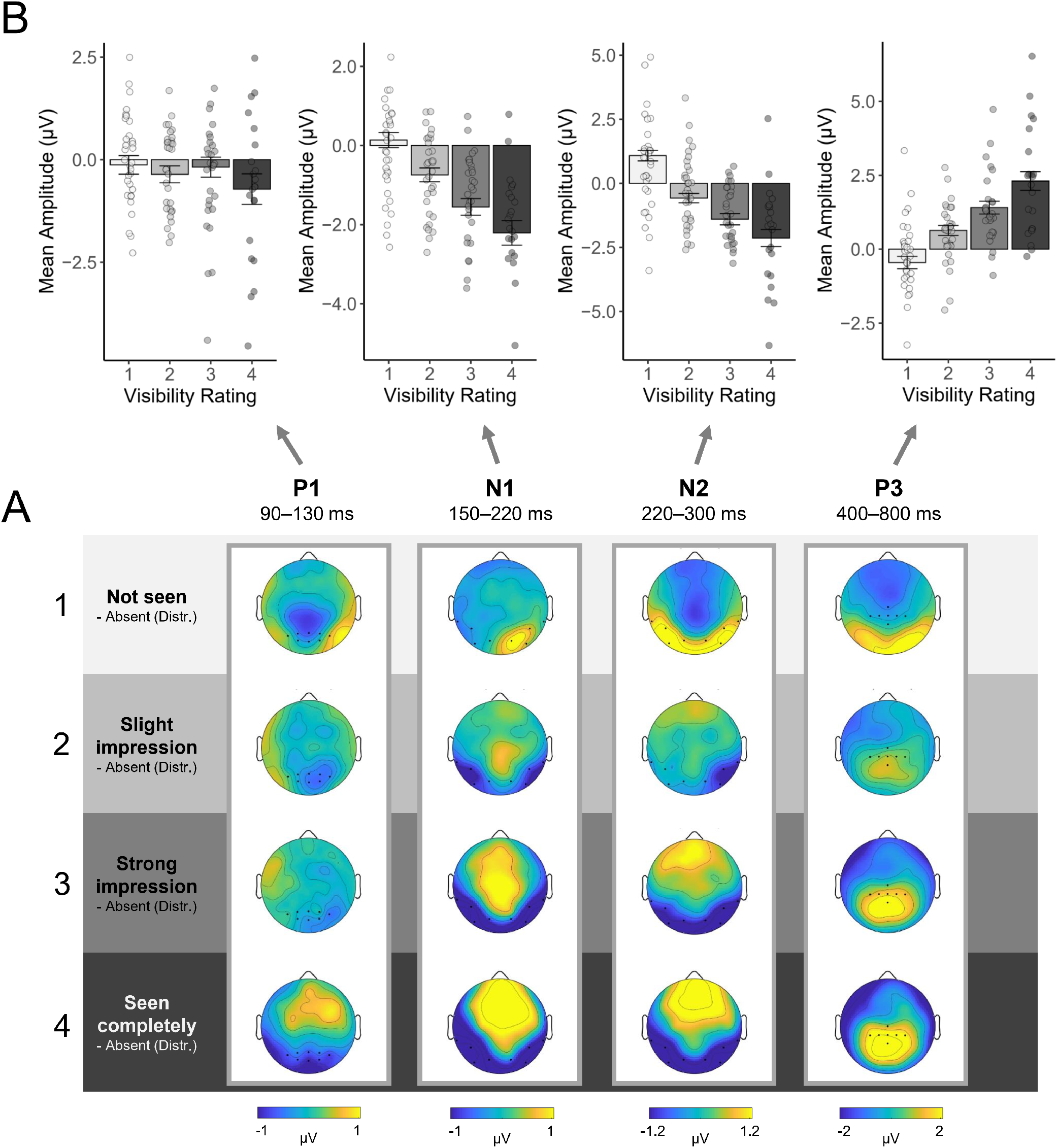
Mean activity during P1, N1, N2 and P3 time range following T2 target onset for each visibility level. **(A)** Topographies show graded differences in the N1, N2, and P3 time range, but not in the P1 time range. Topographies were obtained by subtracting activity during the T2-absent condition (in which a distractor image instead of T2 was shown) in order to remove activity caused by the RSVP stream from relevant activity following T2 onset. Please note that the topographies therefore reflect relative differences to this baseline (distractor only) condition. Due to this subtraction procedure, a reverse pattern of the N1, N2, and P3 component is obtained for *not seen* responses indicating that activity was reduced as compared to a condition where an irrelevant distractor was displayed. Furthermore, since P1 activity was similar for T2 target and distractor stimuli (i.e., a positive deflection for the respective ROI electrodes in both conditions), the subtraction resulted in a mean ROI activity around 0. **(B)** Mean amplitude across respective time range and ROI electrodes for each component, displaying differences in activity between visibility levels (visibility levels: 1 = *not seen*, 2 = *slight impression*, 3 = *strong impression*, 4 = *seen completely*). Amplitudes were obtained by subtracting activity during the T2-absent condition. Error bars depict 95% confidence intervals; circles represent means of individual participants. For illustration of corresponding by-item means, see Figure A2 in the appendix.

***P1***. Comparison of adjacent visibility levels yielded no significant differences during the P1 time range (all *p*s *≥*.099). *≤*

***N1***. The analyses yielded significant differences for all successive comparisons of adjacent visibility levels (all *p*s≤ .012).

***N2***. The analyses yielded significant differences for all successive comparisons of adjacent visibility levels (all *p*s <.001).

***P3***. The analyses yielded significant differences for all successive comparisons of adjacent visibility levels (all *p*s <.001).

## Discussion

The present study examined the form of visual consciousness during the attentional blink with subjective visibility ratings on the PAS scale, an additional objective criterion, and the measurement of ERPs (P1, N1, N2, and P3) time-locked to T2 onset. We observed a high proportion of intermediate scale responses in subjective visibility ratings, an increase in accuracy as well as a decrease in reaction time in the objective gender classification task with increasing visibility ratings, and graded ERP modulations associated with differences in visibility in the N1, N2, and P3 components. Only the earliest tested visual component, the P1, did not show a graded pattern. These findings suggest the presence of graded states of visual consciousness in the attentional blink.

The visibility ratings showed a graded pattern, as previously reported in attentional blink studies by Nieuwenhuis and de Kleijn (2011; when individual letters were used as T2 targets) and Pretorius (2014, Experiment 1), but in contrast to the behavioral pattern reported by Sergent and Dehaene (2004), and Sergent et al. (2005). This difference may be related to the types of stimuli employed. The use of words in the attentional blink may cause a bias towards a dichotomous response pattern because partial awareness of the stimulus (e.g., perceiving only parts of the word) may result in not being able to recognize it as an existing word in the first place and, thus, falsely lead to a “not seen” rating. Faces, on the other hand, may still be recognized as such even in the case of degraded or incomplete perception, e.g., even when more specific information about identity may not be fully available. Furthermore, differences may be due to a higher sensitivity of the four-point PAS scale used here as compared to the 21-point scale used by Sergent et al. (2005), in regard to being able to capture graded states of consciousness (Pretorius et al., 2016).

The combination of the subjective visibility rating with an objective task enabled us to examine systematic differences between visibility levels related to objective performance. As would be expected given graded conscious perception, the accuracy in the objective task increased with increasing levels of subjective visibility. The sharp increase in performance from *not seen* to *slight impression* demonstrates that already a low level of perceived visibility is sufficient to correctly identify the gender in almost 90% of cases—which, however, still improved significantly up to almost 100% in case of a *seen completely* rating. The high performance in the gender task may be due to the very rapid decoding of gender information from faces (Dobs et al., 2019) and to the ability to make judgments about gender also based on individual facial features (Brown & Perrett, 1993). Differences between visibility ratings were furthermore reflected in gradual differences in reaction times for the gender classification task, with the longest reaction times in case of *not seen* ratings and fastest reaction in case of *seen completely* ratings. These differences may be indicative of the ease with which the face’s gender could be determined. They provide additional evidence for the presence of systematic differences between different levels of subjective visibility.

In the ERPs, no differences in P1 activity were found for the comparison of (adjacent) visibility levels, in line with Sergent et al. (2005). Thus, differences in visibility ratings cannot be attributed to differences in low-level visual processing of T2 stimuli. This result is consistent with overall evidence indicating that activity during the P1 time range is not modulated by the attentional blink (see Zivony & Lamy, 2020). In the consciousness literature as a whole, there also exists comparatively little evidence for differences in the P1 time range, and by now, there seems to be a growing consensus that the P1 is not a marker of conscious awareness (Förster et al., 2020; Koivisto & Revonsuo, 2010).

Already in the N1/N170 component, significant gradations of activity depending on the reported visibility were found. This effect stands in contrast to the attentional blink literature in general (when comparing “seen” and “unseen” trials) which overall indicates no differences during this time range (see Zivony & Lamy, 2020). In Sergent et al. (2005), also no differences were observed in the N1 component. Yet, small initial differences were found to occur already during the N1 time range around N170 ms, but at a different location—over central electrodes—in form of a slightly stronger positivity for seen compared to unseen word stimuli. These differential results may be due to differences in the materials used (e.g., differences in the processing of faces as compared to word stimuli, see e.g. Aranda et al., 2010; Maurer et al., 2008), possibly also related to a preferential processing and early selection of faces as a very important source of social information compared to other stimuli (see Darque et al., 2012). Yet, modulations of the N170 in the attentional blink have not always been found for face stimuli either (Harris et al., 2013) and, thus, further variables of the study design (e.g., the SOA between T1 and T2, and the kind of T2 task used) are likely crucial in this regard as well.

Across different paradigms the evidence on the neural correlates of consciousness also indicates differences between seen and unseen stimuli already during the N1 time range as part of the broader visual awareness negativity (VAN) component which spans the time range of the N1 and N2 components, with differences in onset and peak latencies depending on the specific study designs (Förster et al., 2020; Koivisto & Revonsuo, 2010). In line with these findings, our data indicate the presence of a broad, continuous pattern of posterior negativity extending over the period of the N1 and N2, which can, thus, be interpreted in terms of the VAN component.

Significant graded differences for visibility levels were also found in the N2 time range. Differences between detected and undetected stimuli during this time period have frequently been reported in the attentional blink literature (Koivisto & Revonsuo, 2008; Maier & Abdel Rahman, 2018; Sergent et al., 2005; Zivony & Lamy, 2020; yet, this has not always been found, see e.g., Batterink et al., 2012). In terms of functional significance, modulations during this time range are assumed to reflect differences in attentional engagement (Zivony & Lamy, 2020). The graded pattern observed here is consistent with the one reported by Sergent et al. (2005) where graded differences were observed for the comparison of four visibility levels (which were obtained by separating the used 21-point scale into four categories). As outlined in the introduction, based on the overall result pattern, the authors interpreted these differences to reflect pre-conscious processing rather than conscious perception.

A graded pattern as a function of the visibility level was also found in the P3. This is one of the most important differences to Sergent et al. (2005), who found a dichotomous pattern for this component with a separation into seen or not seen states without gradations. Based on the correspondence to the dichotomous pattern in the behavioral data, they concluded that activity during this period may reflect access to consciousness. It is important to note that on the basis of our data (and this is a difficulty in the search for neural correlates of consciousness in general) it is not possible to directly determine which processing stage reflects (the access to) visual consciousness. Yet, gradations in activity of both VAN and P3 component, which could potentially qualify as markers of conscious perception, strongly indicate a graded form of consciousness during the attentional blink in our study.

Evidence from other paradigms (visual masking or low-contrast stimuli) that have recorded electrophysiological measures (EEG or MEG) in combination with visibility/awareness ratings also report modulations during the VAN time range in connection to reported visibility (Andersen et al., 2016; Derda et al., 2019; Koivisto & Grassini, 2016; Tagliabue et al., 2016). Although P3/LP modulations were observed in these studies as well, they were not found as consistently (e.g., only for a high-level but not low-level condition in Derda et al., 2019) and were less predictive of the visual impression (Andersen et al., 2016; Koivisto & Grassini, 2016) as compared to the VAN. Overall, these studies—in line with overall evidence (Förster et al., 2020)—indicate that activity during the VAN time range reflects subjective perceptual experience of a stimulus (rather than pre-conscious processing), whereas modulations in the P3/LP time range may reflect post-perceptual processing, such as further conscious processing of task-relevant features in working memory (Koivisto & Grassini, 2016), accumulation of sensory evidence (Melloni et al., 2011) or the conscious experience of non-perceptual information (Derda et al., 2019). Further studies are needed to determine whether these conclusions can be applied to the attentional blink paradigm as well.

### Limitations and Implications for Further Research

A limitation of the present study with respect to the distribution of visibility ratings is that there was no single task condition (in which participants respond to T2 only, and not to T1) and thus no evidence of what the distribution of visibility ratings looks like under a condition of non-disrupted attention. Such a condition would be useful to compare—as in Sergent et al. (2005)—the visibility distribution in the attentional blink with the theoretical distribution in the case of dichotomous conscious perception, which can be estimated by the combination of unseen states (when T2 is absent in a single task condition) and seen states (when T2 is present in a single task condition). The long lag condition is only of limited informative value here since the (objective) performance is significantly better, but not at ceiling level. The comparison shows a typical attentional blink pattern with reduced detection in short lag trials. However, to ensure that the degraded T2 detection can be clearly attributed to the limitations of processing resources caused by the dual task and is not also generally limited by a high task difficulty (e.g., due to high visual demands), subsequent studies should include a single task condition. At the same time, the present data does indicate that (assumedly) complete perception is possible and does occur (in case of a *seen completely* rating) under the present task conditions. The presence of intermediate scale responses therefore should not be attributable merely to a general visual task difficulty limiting how well stimuli can be perceived since such a general limitation would be expected to apply to all trials.

One further methodological limitation of the present study consists in the fact that a relatively small number of *seen completely* trials (total of 938, with an average of 29 trials per participant, range = 0-177) entered the analysis compared to the other levels of visibility. Participants differed in their response behavior, and even though the majority used all four levels of visibility (see Figure A1 in the appendix for individual displays), ten participants had no trials at all for *seen completely* in short lag trials. In addition, participants differed considerably in how many trials per visibility level are included in the analysis (this is a common problem in studies investigating ERPs using the PAS or other visibility scales). However, these differences are taken into account in the LMM analysis, which is based on single trial data, and the results therefore cannot be attributed to distortions in an averaging procedure (due to differences in trial numbers per condition). Furthermore, even if only the first three visibility levels were taken into account, the same conclusions in favor of graded visual consciousness would be reached.

An interesting question which remains open is how exactly degraded perception is manifested. Which aspects of the visual representation exactly are affected by the attentional blink? For example, degradation could refer to a less intense or a less clear percept but also to an only partially perceived stimulus (see, e.g,. Fazekas & Overgaard, 2018 for a differentiation of relevant aspects). Investigating this question may provide deeper insight into the exact mechanisms of the attentional blink.

## Conclusions

The present ERP study provides evidence for graded visual consciousness during the attentional blink. The findings indicate that the specific study design and variations of the stimulus materials may determine whether graded levels of consciousness can be detected. Concerning the use of the attentional blink, as yet, many studies have only taken into account two dichotomous states of “seen” versus “not seen” or “detected” versus “not detected” in order to, for example, test the effects of an experimental manipulation (e.g., emotional influences on access to consciousness). However, considering that visual perception in the attentional blink may not necessarily be dichotomous has important consequences for the study design and the respective conclusions.

## Data and Code Availability

Data and analysis scripts are available to reviewers, and will be made publicly available upon peer-reviewed publication.

## Acknowledgements

We thank Guido Kiecker for technical support, and Friedrich Eiserbeck, Franziska Glogau, Maja Heitmann, Hannah Kaube, Kirsten Stark, and Nura Völk for their help with data collection. This pre-print was created using the Henriques-Lab bioRxiv template.

## APPENDIX

### Tables

**Table A1.** Differences between lags in accuracy in objective task (GLMM statistics) and mean visibility (LMM statistics)

| Predictors | Accuracy |  |  |  | Mean visibility |  |  |  |  |
| --- | --- | --- | --- | --- | --- | --- | --- | --- | --- |
|  | Odds Ratios | CI | *z* | *p* | *b* | CI | *df* | *t* | *p* |
| (Intercept) | 4.53 | 3.17; 6.49 | 8.26 | **<0.001** | 2.38 | 2.18; 2.59 | 36.55 | 22.90 | **<0.001** |
| Lag2-1 | 2.67 | 1.94; 3.69 | 5.99 | **<0.001** | 0.32 | 0.18; 0.45 | 31.00 | 4.67 | **<0.001** |
| Random Effects |  |  |  |  |  |  |  |  |  |
| $\sigma^2$ | 3.29 | | | | 0.53 | | | | |
| $\tau_{00}$ | 0.94 Participant | | | | 0.32 Participant | | | | |
|  | 0.09 Item |  |  |  | 0.02 Item |  |  |  |  |
| $\tau_{11}$ | 0.76 Participant.lag2-1 | | | | 0.14 Participant.lag2-1 | | | | |
|  | 0.01 Item.lag2-1 |  |  |  |  |  |  |  |  |
| $\rho_{01}$ | -0.06 Participant | | | | 0.06 Participant | | | | |
|  | 0.15 Item |  |  |  |  |  |  |  |  |
| ICC | 0.27 |  |  |  | 0.42 |  |  |  |  |
| N | 32 Participant |  |  |  | 32 Participant |  |  |  |  |
|  | 24 Item |  |  |  | 24 Item |  |  |  |  |
| Observations | 16757 |  |  |  | 16757 |  |  |  |  |
| Marginal R2 / Conditional R2 | 0.051 / 0.308 |  |  |  | 0.027 / 0.432 |  |  |  |  |

Note. CI = 95% confidence interval.

**Table A2.** Differences between visibility levels for prediction of accuracy (GLMM statistics) and log-transformed reaction time (LMM statistics) in objective task

| Predictors | Accuracy |  |  |  | Reaction time |  |  |  |  |
| --- | --- | --- | --- | --- | --- | --- | --- | --- | --- |
|  | Odds Ratios | CI | *z* | *p* | *b* | CI | *df* | *t* | *p* |
| (Intercept) | 9.27 | 5.66; 15.17 | 8.85 | **<0.001** | 6.22 | 6.14; 6.30 | 33.38 | 155.36 | **<0.001** |
| Visibility2-1 | 511.86 | 356.81; 734.29 | 33.88 | **<0.001** | -0.04 | -0.07; -0.01 | 7518.70 | -2.75 | **0.006** |
| Visibility3-2 | 8.24 | 5.85; 11.60 | 12.08 | **<0.001** | -0.11 | -0.14; -0.08 | 7477.14 | -7.55 | **<0.001** |
| Visibility4-3 | 10.46 | 2.55; 42.98 | 3.26 | **.001** | -0.09 | -0.13; -0.05 | 7523.08 | -4.49 | **<0.001** |
| Random Effects |  |  |  |  |  |  |  |  |  |
| $\sigma^2$ | 3.29 | | | | 0.21 | | | | |
| $\tau_{00}$ | 0.93 Participant | | | | 0.05 Participant | | | | |
|  | 0.00 Item |  |  |  | 0.00 Item |  |  |  |  |
| ICC | 0.22 |  |  |  | 0.19 |  |  |  |  |
| N | 32 Participant |  |  |  | 32 Participant |  |  |  |  |
|  | 24 Item |  |  |  | 24 Item |  |  |  |  |
| Observations | 8540 |  |  |  | 7530 |  |  |  |  |
| Marginal R2 / Conditional R2 | 0.756 / 0.810 |  |  |  | 0.025 / 0.21 |  |  |  |  |

Note. CI = 95% confidence interval.

## APPENDIX

### Figures

**Fig. A1.**
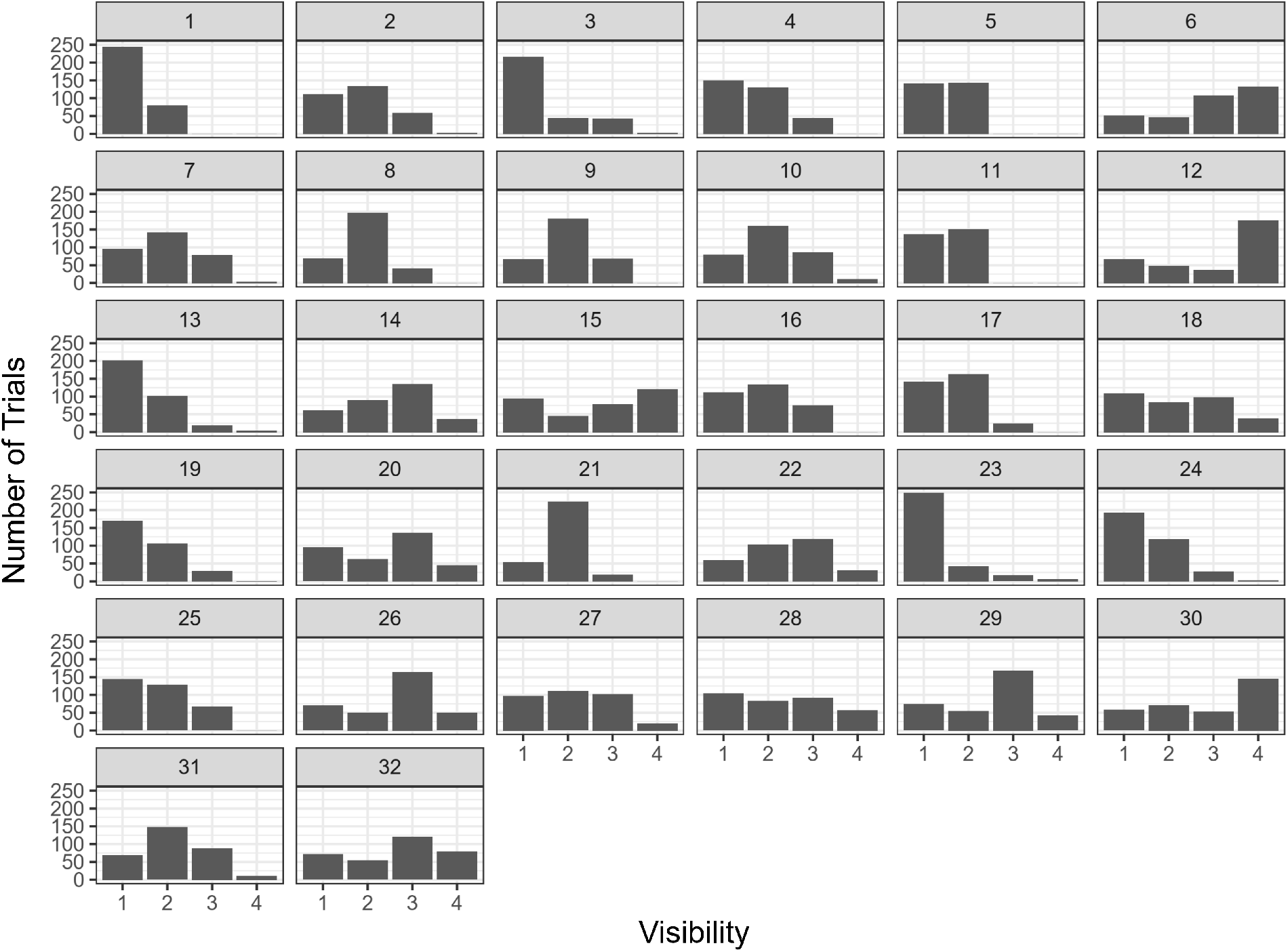
Distribution of visibility ratings in T2 present short lag trials for individual participants. Visibility levels: 1 = *not seen*, 2 = *slight impression*, 3 = *strong impression*, 4 = *seen completely*.

**Fig. A2.**
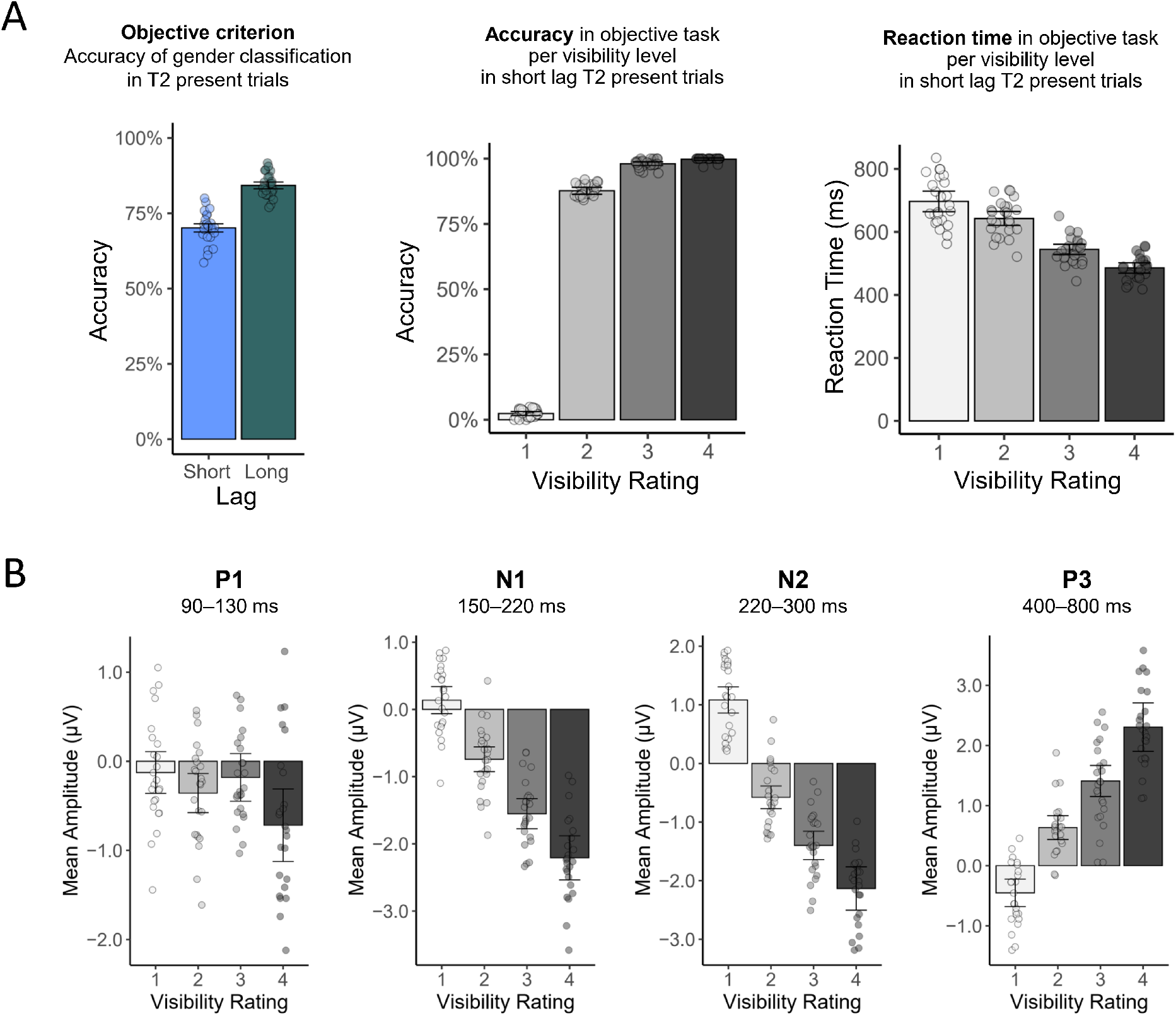
Illustration of results including by-item means depicted as circles. **(A)** Behavioral results. **(B)** ROI activity in short lag T2-present trials. Amplitudes were obtained by subtracting activity during the T2-absent condition (in which a distractor image instead of T2 was shown) in order to remove activity caused by the RSVP stream from relevant activity following T2 onset. Error bars represent 95% confidence intervals. Visibility levels: 1 = *not seen*, 2 = *slight impression*, 3 = *strong impression*, 4 = *seen completely*.

## Footnotes

1 Further, the measure—post-decisional wagering (PDW) instead of visibility ratings—was found to influence the pattern of results, leading to a uni-modal pattern in the case of words as stimuli and to a graded pattern in the case of single numbers. However, it needs to be considered that the measure of PDW does not directly refer to the perceived visibility or quality of the percept but instead depends on a metacognitive evaluation of the precision of a judgement. The pattern of results might thereby be influenced by additional factors such as risk aversion (for a comparison of different awareness scales, see Wierzchon’ et al., 2014).

2 In a further EEG study (Pincham et al. 2016), subjective visibility of T2 letter targets was measured on a 6-point scale in addition to assessing T2 accuracy. The results showed that T2 accuracy and subjective visibility are distinguishable (especially in early lags) and that P3 activity is more strongly associated with visibility differences. However, to analyze ERPs corresponding to visibility differences, the visibility responses were summarized in two bins (the two lowest ratings as “low visibility” and the four highest as “high visibility”) in order to obtain comparable trial numbers for the ERP analysis. Thus, no gradual differences were tested in the ERPs. Furthermore, no modulation in earlier components (N2/VAN) was reported.

3 Please note that, as yet, there seems to be uncertainty as to which scale type the PAS can be considered as (interval or ordinal scale, see Sandberg Overgaard, 2015, p. 192) and, thus, whether the requirements for linear mixed models—measurement of the dependent variable on an interval or ratio scale—are fulfilled. However, even if this should not be the case, it has been shown that LMMs are usually robust in case of ordinal scale level (see, e.g., Norman, 2010).

## References

Andersen, L. M., Pedersen, M. N., Sandberg, K., & Overgaard, M. (2016). Occipital MEG activity in the early time range (<300 ms) predicts graded changes in Perceptual consciousness. Cerebral Cortex, 26(6), 2677–2688. https://doi.org/10.1093/cercor/bhv108

Aranda, C., Madrid, E., Tudela, P., & Ruz, M. (2010). Category expectations: A differential modulation of the N170 potential for faces and words. Neuropsychologia, 48(14), 4038–4045. https://doi.org/10.1016/j.neuropsychologia.2010.10.002

Asplund, C. L., Fougnie, D., Zughni, S., Martin, J. W., & Marois,R. (2014). The attentional blink reveals the probabilistic nature of discrete conscious perception. Psychological Science, 25(3), 824–831. https://doi.org/10.1177/0956797613513810

Bates, D., Kliegl, R., Vasishth, S., & Baayen, H. (2018). Parsimonious Mixed Models. http://arxiv.org/abs/1506.04967

Bates, D., Mächler, M., Bolker, B. M., & Walker, S. V. (2015). Fitting linear mixed-effects models using lme4. Journal of Statistical Software, 67(1). https://doi.org/10.18637/jss.v067.i01

Batterink, L., Karns, C. M., & Neville, H. (2012). Dissociable mechanisms supporting awareness: The P300 and gamma in a linguistic attentional blink task. Cerebral Cortex, 22(12), 2733–2744. https://doi.org/10.1093/cercor/bhr346

Blumenfeld, H. (2014). A master switch for consciousness? Epilepsy & Behavior, 37, 234–235. https://doi.org/10.1016/j.yebeh.2014.07.008

Brown, E., & Perrett, D. I. (1993). What gives a face its gender? Perception, 22, 829–840. https://doi.org/10.1068/p220829

Cohen, M. A., Ortego, K., Kyroudis, A., & Pitts, M. (2020). Distinguishing the neural correlates of perceptual awareness and postperceptual processing. The Journal of Neuroscience, 40(25), 4925–4935. https://doi.org/10.1523/jneurosci.0120-20.2020

Darque, A., Del Zotto, M., Khateb, A., & Pegna, A. J. (2012). Attentional modulation of early ERP components in response to faces: Evidence from the attentional blink paradigm. Brain Topography, 25(2), 167–181. https://doi.org/10.1007/s10548-011-0199-5

Dehaene, S., Sergent, C., & Changeux, J. P. (2003). A neuronal network model linking subjective reports and objective physiological data during conscious perception. Proceedings of the National Academy of Sciences of the United States of America, 100(14), 8520–8525. https://doi.org/10.1073/pnas.1332574100

Del Cul, A., Baillet, S., & Dehaene, S. (2007). Brain dynamics underlying the nonlinear threshold for access to consciousness. PLoS Biology, 5(10), 2408–2423. https://doi.org/10.1371/journal.pbio.0050260

Delorme, A., & Makeig, S. (2004). EEGLAB: an open source toolbox for analysis of single-trial EEG dynamics. Journal of Neuroscience Methods, 13, 9–21. https://doi.org/http://dx.doi.org/10.1016/j.jneumeth.2003.10.009

Derda, M., Koculak, M., Windey, B., Gociewicz, K., Wierzchon, M., Cleeremans, A., & Binder, M. (2019). The role of levels of processing in disentangling the ERP signatures of conscious visual processing. Consciousness and Cognition, 73, 102767. https://doi.org/10.1016/j.concog.2019.102767

Dobs, K., Isik, L., Pantazis, D., & Kanwisher, N. (2019). How face perception unfolds over time. Nature Communications, 10(1), 1–10. https://doi.org/10.1038/s41467-019-09239-1

Eiserbeck, A., Enge, A., Rabovsky, M., & Abdel Rahman, R. (2021). (Dis)Trust before first sight: Knowledge- and appearance-based trustworthiness effects on visual consciousness of faces. Manuscript in preparation.

Elliott, J. C., Baird, B., & Giesbrecht, B. (2016). Consciousness isn’t all-or-none: Evidence for partial awareness during the attentional blink. Consciousness and Cognition, 40, 79–85. https://doi.org/10.1016/j.concog.2015.12.003

Fazekas, P., & Overgaard, M. (2018). A multi-factor account of degrees of awareness. Cognitive Science, 42(6), 1833–1859. https://doi.org/10.1111/cogs.12478

Förster, J., Koivisto, M., & Revonsuo, A. (2020). ERP and MEG correlates of visual consciousness: The second decade. Consciousness and Cognition, 80, 102917. https://doi.org/10.1016/j.concog.2020.102917

Frömer, R., Maier, M., & Abdel Rahman, R. (2018). Group-level EEG-processing pipeline for flexible single trial-based analyses including linear mixed models. Frontiers in Neuroscience, 12, 1–15. https://doi.org/10.3389/fnins.2018.00048

Harris, J. A., McMahon, A. R., & Woldorff, M. G. (2013). Disruption of visual awareness during the attentional blink is reflected by selective disruption of late-stage neural processing. Journal of Cognitive Neuroscience, 25(11), 1863–1874. https://doi.org/10.1162/jocna00443

Hinojosa, J. A., Mercado, F., & Carretié, L. (2015). N170 sensitivity to facial expression: A meta-analysis. Neuroscience and Biobehavioral Reviews, 55, 498–509. https://doi.org/10.1016/j.neubiorev.2015.06.002

Ille, N., Berg, P., & Scherg, M. (2002). Artifact correction of the ongoing EEG using spatial filters based on artifact and brain signal topographies. Journal of Clinical Neurophysiology, 19(2), 113–124. https://doi.org/10.1097/00004691-200203000-00002

Itier, R. J., & Taylor, M. J. (2004). N170 or N1? Spatiotemporal differences between object and face processing using ERPs. Cerebral Cortex, 14(2), 132–142. https://doi.org/10.1093/cercor/bhg111

Jonkisz, J., Wierzchon, M., & Binder, M. (2017). Four-dimensional graded consciousness. Frontiers in Psychology, 8(MAR), 1–10. https://doi.org/10.3389/fpsyg.2017.00420

Kiefer, M., & Kammer, T. (2017). The emergence of visual awareness: Temporal dynamics in relation to task and mask type. Frontiers in Psychology, 8, 1–14. https://doi.org/10.3389/fpsyg.2017.00315

Koch, C., Massimini, M., Boly, M., & Tononi, G. (2016). Neural correlates of consciousness: Progress and problems. Nature Reviews Neuroscience, 17(5), 307–321.https://doi.org/10.1038/nrn.2016.22

Koivisto, M., & Grassini, S. (2016). Neural processing around 200 ms after stimulus-onset correlates with subjective visual awareness. Neuropsychologia, 84, 235–243. https://doi.org/10.1016/j.neuropsychologia.2016.02.024

Koivisto, M., & Revonsuo, A. (2008). Comparison of event-related potentials in attentional blink and repetition blindness. Brain Research, 1189(1), 115–126. https://doi.org/10.1016/j.brainres.2007.10.082

Koivisto, M., & Revonsuo, A. (2010). Event-related brain potential correlates of visual awareness. Neuroscience and Biobehavioral Reviews, 34(6), 922–934. https://doi.org/10.1016/j.neubiorev.2009.12.002

Kuznetsova, A., Brockhoff, P. B., & Christensen, R. H. B. (2017). lmerTest package: Tests in linear mixed effects models. Journal of Statistical Software, 82(13). https://doi.org/10.18637/jss.v082.i13

Ma, D. S., Correll, J., & Wittenbrink, B. (2015). The Chicago face database: A free stimulus set of faces and norming data. Behavior Research Methods, 47(4), 1122–1135. https://doi.org/10.3758/s13428-014-0532-5

Maier, M., & Abdel Rahman, R. (2018). Native language promotes access to visual consciousness. Psychological Science, 29(11), 1757–1772. https://doi.org/10.1177/0956797618782181

Maurer, U., Rossion, B., & McCandliss, B. D. (2008). Category specificity in early perception: Face and word N170 responses differ in both lateralization and habituation properties. Frontiers in Human Neuroscience, 2(18), 1–7. https://doi.org/10.3389/neuro.09.018.2008

Melloni, L., Schwiedrzik, C. M., Müller, N., Rodriguez, E., & Singer, W. (2011). Expectations change the signatures and timing of electrophysiological correlates of perceptual awareness. Journal of Neuroscience, 31(4), 1386–1396. https://doi.org/10.1523/JNEUROSCI.4570-10.2011

Müsch, K., Engel, A. K., & Schneider, T. R. (2012). On the blink: The importance of target-distractor similarity in eliciting an attentional blink with faces. PLoS ONE, 7(7). https://doi.org/10.1371/journal.pone.0041257

Nieuwenhuis, S., & de Kleijn, R. (2011). Consciousness of targets during the attentional blink: A gradual or all-or-none dimension? Attention, Perception, and Psychophysics, 73(2), 364–373. https://doi.org/10.3758/s13414-010-0026-1

Norman, G. (2010). Likert scales, levels of measurement and the “laws” of statistics. Advances in Health Sciences Education, 15(5), 625–632. https://doi.org/10.1007/s10459-010-9222-y

Overgaard, M., Rote, J., Mouridsen, K., & Ramsøy, T. Z. (2006). Is conscious perception gradual or dichotomous? A comparison of report methodologies during a visual task. Consciousness and Cognition, 15(4), 700–708. https://doi.org/10.1016/j.concog.2006.04.002

Pincham, H. L., Bowman, H., & Szucs, D. (2016). The experiential blink: Mapping the cost of working memory encoding onto conscious perception in the attentional blink. Cortex, 81, 35–49. https://doi.org/10.1016/j.cortex.2016.04.007

Pitts, M. A., Metzler, S., & Hillyard, S. A. (2014). Isolating neural correlates of conscious perception from neural correlates of reporting one’s perception. Frontiers in Psychology, 5, 1–16. https://doi.org/10.3389/fpsyg.2014.01078

Pretorius, H. (2014). Is conscious perception a continuous or dichotomous phenomenon? [Doctoral dissertation, University of Cape Town]. https://open.uct.ac.za/bitstream/handle/11427/12965/thesis_hum_2014_pretorius_h.pdf?sequence=1

Pretorius, H., Tredoux, C., & Malcolm-Smith, S. (2016). Subjective awareness scale length influences the prevalence, not the presence, of graded conscious states. Consciousness and Cognition, 45, 47–59. https://doi.org/10.1016/j.concog.2016.08.007

R Core Team. (2014). R: A language and environment for statistical computing. R Foundation for Statistical Computing. https://www.r-project.org/

Ramsøy, T. Z., & Overgaard, M. (2004). Introspection and perception. Phenomenology and the Cognitive Sciences, 3(1), 1–23. https://doi.org/10.1007/BF00776206

Raymond, J. E., Shapiro, K. L., & Arnell, K. M. (1992). Temporary suppression of visual processing in an RSVP task: An attentional blink? In Journal of Experimental Psychology: Human Perception and Performance (Vol. 18, Issue 3, pp. 849–860). https://doi.org/10.1037/0096-1523.18.3.849

Sandberg, K., & Overgaard, M. (2015). Using the perceptual awareness scale (PAS). In Behavioural methods in consciousness research (pp. 181–195). https://doi.org/10.1093/acprof:oso/9780199688890.001.0001

Sergent, C., Baillet, S., & Dehaene, S. (2005). Timing of the brain events underlying access to consciousness during the attentional blink. Nature Neuroscience, 8(10), 1391–1400. https://doi.org/10.1038/nn1549

Sergent, C., & Dehaene, S. (2004). Is consciousness a gradual phenomenon? Psychological Science, 15(11), 720–728. https://doi.org/10.1111/j.0956-7976.2004.00748.x

Stein, T., & Peelen, M. V. (2020). Dissociating conscious and unconscious influences on visual detection effects. Nature Human Behaviour. https://doi.org/10.1038/s41562-020-01004-5

Tagliabue, C. F., Mazzi, C., Bagattini, C., & Savazzi, S. (2016). Early local activity in temporal areas reflects graded content of visual perception. Frontiers in Psychology, 7, 1–10. https://doi.org/10.3389/fpsyg.2016.00572

Wierzchon, M., Paulewicz, B., Asanowicz, D., Timmermans, B., & Cleeremans, A. (2014). Different subjective awareness measures demonstrate the influence of visual identification on perceptual awareness ratings. Consciousness and Cognition, 27(1), 109–120. https://doi.org/10.1016/j.concog.2014.04.009

Willenbockel, V., Sadr, J., Fiset, D., Horne, G. O., Gosselin, F., & Tanaka, J. W. (2010). Controlling low-level image properties: The SHINE toolbox. Behavior Research Methods, 42(3), 671–684. https://doi.org/10.3758/BRM.42.3.671

Windey, B., Vermeiren, A., Atas, A., & Cleeremans, A. (2014). The graded and dichotomous nature of visual awareness. Philosophical Transactions of the Royal Society B: Biological Sciences, 369(1641). https://doi.org/10.1098/rstb.2013.0282

Zivony, A., & Lamy, D. (2020). What processes are disrupted during the attentional blink? An integrative review of event-related potentials research. https://doi.org/10.31234/osf.io/epfbt

